# State Space Misspecification in Morphological Phylogenetics: A Pitfall for Models and Parsimony Alike

**DOI:** 10.1101/2025.04.22.650124

**Authors:** EJ Huang

## Abstract

Phylogenetic analysis relies on two fundamental levels of biological information: genotype and phenotype. Molecular data benefit from operating within a well-defined, finite state space (e.g., nucleotide alphabets), whereas morphological data present inherent challenges due to frequently ambiguous character states and variable state counts. In this study, I use simulated data to examine how state space misspecification (SSM), defined as the mismatch between the assumed and true state space, affects phylogenetic reconstruction. Results show that SSM generally reduces topological accuracy, with the extent of its impact depending on mutation rate, state space disparity, and the proportion of affected characters. Counterintuitively, under conditions typical of empirical morphological datasets (high proportions of binary characters and elevated mutation rates), SSM can improve topological precision. This creates a paradox where an incorrect model outperforms a correct one, though at the cost of distorted branch lengths. Importantly, the effects of SSM extend beyond model-based approaches. I demonstrate, through an extension of the no common mechanism (NCM) model, that standard maximum parsimony is consistent with the assumption that characters evolved under an SSM model—a largely overlooked feature. To address this, I propose a state-space-aware weighting scheme that accounts for variation in character state space. I also discuss additional strategies for mitigating SSM, including model adjustments and reducing reliance on oversimplified binary coding. This work underscores the need to explicitly address state space uncertainty in morphological phylogenetics. As morphology remains crucial for reconstructing deep-time lineages and integrating fossils, accounting for SSM is essential to improving the reliability of evolutionary trees.

## Introduction

Our ability to accurately and precisely infer the evolutionary relationships of organisms relies on robust phylogenetic methods. The primary source of phylogenetic inference is homology, or the information shared among lineages due to common descent (Hennig 1966; de Pinna 1991; Nixon and Carpenter 2012). Homology is assessed at two levels of biological information. One level is the genotype, usually analyzed in the form of nucleotide or amnio acid sequences, as well as via other less-utilized molecules (Brown 2002; Briggs and Summons 2014). For instance, the first molecular phylogenies were assembled not from DNA or protein sequences, but rather from the results of immunological experiments (Nuttall et al. 1904; Brown 2002). In the modern research context, rapid advances in sequencing technology have provided workers with efficient and inexpensive techniques (Satam et al. 2023) capable of generating enormous amounts of molecular data for use in phylogenetic analysis (Benson et al. 2013). Additionally, changes in nucleotide and amino acid sequences are rather limited. Each locus contains only one of four possible nucleotides and only one of around twenty possible amino acids. This highly constrained paradigm of substitution is the foundation on which most modern phylogenetic methods are built (Goldman 1993; Huelsenbeck and Crandall 1997; Huelsenbeck et al. 2001; Arenas, 2015; Nascimento et al. 2017). The paradigm benefits from little ambiguity in defining characters, character states, or in recognizing the size of state space (Wagner 2000; Cuthill 2015). Here, I define state space as the set of all possible states for a given character.

The other level is the phenotype, mostly analyzed in the form of morphological matrices. Early phylogenetic studies (Ragan 2009) necessarily depended on phenotypic data, as they predated both the discovery of DNA (Lamm et al. 2020) and the development of protein/DNA sequencing techniques (Brown 2002). However, morphology-based phylogenetics presents a number of unique challenges (Wiens, 2001; Parins-Fukuchi 2018). The challenges stem from the inherently time-consuming and conceptually fraught task of partitioning what are often complex developmental products into modular units (i.e., characters) and their meaningful variations (i.e., character states). Consequently, not only can state definitions be subject to scrutiny, but also the true state spaces of many characters are unknown (Donoghue and Ree 2000; Cuthill 2015). That is, even if biologists carefully lay out the observed state space for each analyzed character, there is often little reason to assume that such descriptions exhaust the possibility of variation. Morphological character state space has been studied from various perspectives (e.g., Donoghue and Ree 2000; Wiens 2001; Cuthill 2015; Gerber 2018), but a topic that is surprisingly understudied is the consequence of state space being misspecified (e.g., when binary characters are treated as multistate characters) when performing a model-based phylogenetic reconstruction (Casali et al. 2023; Mulvey et al. 2025).

This lack of understanding is crucial because many studies using maximum likelihood estimation only have a single substitution model for morphological characters (e.g., Berger et al. 2011; Kuntner 2013; Scarpetta 2020; Francisco et al. 2024; Torres et al. 2024). The choice of a single substitution model likely reflects the default setting in RAxML (Stamatakis 2014) and IQ-TREE (Nguyen et al. 2015), two of the commonly used likelihood-based phylogenetic inference programs, despite options for multiple substitution models. With a single substitution model, all characters are treated as though they can potentially take on *r* states, where *r* is the size of the observed maximum state space. Since binary and ternary characters dominate most morphological matrices, a single character with large state space would lead to incorrect state space allocated to most characters. In contrast, commonly used Bayesian phylogenetic programs, including MrBayes (Huelsenbeck et al. 2001) and Beast2 (Bouckaert et al. 2014), parse out characters with different observed state spaces by default. Characters with different state space sizes are then assigned different substitution models. Variation noted between reconstructed phylogenies with these programs can thus be a result of differential treatment of state space in addition to the methods themselves.

Besides maximum likelihood estimation and Bayesian inference, maximum parsimony (MP) is not entirely immune to state space uncertainty. Morphological phylogenetics has long relied on maximum parsimony, despite criticism that the approach is prone to long branch attraction, among other issues (Felsenstein 1978, Felsenstein 1983, Scottland and Steel 2015; O’Reilly 2016). A major concern is that unlike molecular data (where sites can be more easily conceptualized as random draws from a population), each morphological character often represents a unique and internally complex biological system. Thus, the use of a model-based approach to describe such characters can be discouraging, especially when it comes to defining model parameters.

On the other hand, although MP is typically considered a non-model-based approach, Tuffley and Steel (1997) demonstrated that under the parameter-abundant ‘no common mechanism’ model (NCM), a topology estimated with maximum likelihood is guaranteed to be identical to one that is produced by maximum parsimony criterion. In fact, NCM has been used to justify the practice of maximum parsimony over other model-based approaches (Farris 2008; Goloboff et al. 2019). However, one assumption of NCM has been largely overlooked: for NCM to consistently select the most parsimonious topology, all characters must assume the same state space size. Specifically, the ML estimate under NCM is given by:

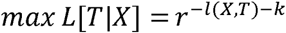

where *r* is the number of possible states and *l(X,T)* is the number of steps required for topology T given character matrix X. While the connection between the two methods does not mean MP explicitly assumes equal state space (Sober 2004), it can indicate topological preference. Thus, whilst commonly described as being ‘model free’, it can be argued that maximum parsimony does in fact make assumptions about character state-space a priori.

In this study, I simulated discrete character data and performed phylogenetic reconstruction to study the effect of state-space misspecification under various scenarios. I also tested how state-space misspecification can affect empirical studies by applying different models to the two published matrices out of our lab (Bever et al. 2015; Torres et al. 2024). These results provide critical assessment of the relationship between state-space misspecification and various parameters of evolutionary models, as well as their impact on overall reconstruction precision. This study represents an important step in improving the ability of morphological phylogenetics to reliably reconstruct evolutionary relationships, and will be most valuable to researchers studying the deep origins of modern biodiversity through morphological variation amongst fossil taxa.

## Materials & Methods

### Data simulation

I generated discrete numerical matrices and performed analyses using R, along with the following packages: ape v5.7.1 (Paradis and Schliep 2019), Matrix v1.5.4.1 (Bates et al. 2010), phytools v1.5.1 (Revell 2012), TreeDist v2.6.3 (Smith 2022), phangorn (Schliep 2011), process (Csárdi and Chang 2025), doParallel (Microsoft and Weston 2022a), foreach (Microsoft and Weston 2022b), and readr v2.1.4 (Wickham et al. 2023). The R script can be found on GitHub (https://github.com/ej91016/SSM). For all but one scenario (see Scenarios for the exception) described in the following paragraphs and Table 1, I first generated 50 trees under pure-birth process (Yule 1925) with a birth rate of one and rescaled all to a tree height of one. I then generated 30 discrete numerical matrices for each tree under the MK model (Lewis 2001) using the ‘sim.Mk’ function in phytools. The parameters modified for different scenarios include the tree size, matrix size, state space size, and mutation rate. The mutation rate is defined as the expected number of mutations per site per unit time.

**Table 1.** Parameter setups for simulation scenarios. For false-space (FS) simulations, the sizes of the modeled state space are listed under multi-state. The parameter(s) modified for each scenario is represented as a function of ‘x’, which is an integer and ranges from 1 to 10, except for the “state space size” scenario where x is an integer and ranges from 2 to 10, “FS d” where it ranges from 1 to 20, and “binary proportion” where it ranges from 0 to 10.

| Scenario | Number of tips | Partition A |  | Partition B |  | Rate | Results |
| --- | --- | --- | --- | --- | --- | --- | --- |
|  |  | State space size | Quantity | State space size | Quantity |  |  |
| Control | 16 | 2 | 500 | 2 | 500 | 0.1x | Figure 1 |
| State space size | 16 | 2 | 500 | x | 500 | 5 |  |
| Tree size | 8x | 2 | 500 | 7 | 500 | 0.5 |  |
| Matrix size | 16 | 2 | 50x | 7 | 50x | 0.5 |  |
| Rate & proportion a | 16 | 2 | 100 | 7 | 900 | 0.1x | Figure 2 |
| Rate & proportion b | 16 | 2 | 500 | 7 | 500 | 0.1x |  |
| Rate & proportion c | 16 | 2 | 900 | 7 | 100 | 0.1x |  |
| Rate & proportion d | 16 | 3 | 900 | 7 | 100 | 0.1x |  |
| Binary proportion | 16 | 2 | 100x | 7 | 100*(10-x) | 0.5 |  |
| False-space a | 16 | 2 | 1000 | 7 | 0 | 0.1x | Figure 3 |
| False-space b | 16 | 3 | 1000 | 7 | 0 | 0.1x |  |
| False-space c | 16 | 3 | 1000 | 8 | 0 | 0.1x |  |
| False-space d | 32 | 2 | 1000 | 7 | 0 | 0.1x | Figure 4,5 |

Each of these matrices consists of two partitions, one binary and the other n-state, where n is the assigned state space. For each of the simulated matrices, I performed maximum likelihood phylogenetic reconstruction using RAxML version 8.2.10 (Stamatakis 2016) with the -m MULTICAT, -K MK, and -V arguments to ensure a MK model without rate heterogeneity, although I did not use the -F option which will disable the final gamma optimization. There was no correction for ascertainment bias as these matrices include invariant characters. For the state-space-aware setup (SSA), a binary MK model was specified for the binary characters and a k-state MK model was specified for characters with a state space size of n. For the state-space-misspecified setup (SSM), both binary characters and k-state characters were assigned to an n-state MK model. Maximum parsimony (MP) trees, when included, are reconstructed using PAUP* v4.0a169 (Swofford 2003) with the TBR swapping algorithm. Each reconstruction involves 20 random starting trees and search for 10,000,000 rearrangements, with a reconnection limit of 20 and allowance to swap on all trees. The parsimony analysis was designed such that the first tree was saved for further analysis, although I noted that there is only one most parsimonious tree for all simulations that were randomly selected to check for script behavior.

I used TreeDist to compute tree distances between the reconstructed phylogenies and the phylogenies from which the matrices were generated. Given the variation and the progression of proposed tree distance measurements, here I use Robinson-Foulds distance (RFD), phylogenetic information distance (PID), and clustering information distance (CID) to incorporate both classic and recent approaches (Robinson and Fould 1981; Smith 2020). For RFD, distances are measured in terms of number of splits. For PID and CID, distances are measured in bits and normalized using the phylogenetic and clustering entropy of the reference tree. In the results and discussion sections, I report primarily CID unless otherwise specified, as it is the least prone to unwanted influences, in particular tree size (Smith 2020). Finally, the average tree distances of the 1500 matrices (50 trees * 30 matrices per tree) under SSA and SSM are plotted for comparison across different parameter values regarding phylogenetic reconstruction precision.

### Scenarios

The scenarios and the corresponding parameter values are listed in Table 1. I first set up a control analysis where the two partitions of the matrices both consist of binary characters. This is to ensure that partition on its own would have no impact on the results. I next tested the effect of differences in state space size by comparing the reconstruction precision when the second partition consists of multistate characters ranging from two to ten states. Based on the result of this analysis (see Results and Discussion), I fixed the state space size of the second partition to seven to reduce the complexity of scenarios in subsequent simulations. After testing the impact of tree size on tree reconstruction with multiples of eight, I also fixed the tree size to 16 for the remaining simulations. The impact of sample size on SSM was tested by setting the total number of characters to multiples of 100, ranging from 100 to 1000. For both tree size and matrix size, I set the proportion of binary characters to 50% and mutation rate to 0.5.

For mutation rate and binary character proportion, I performed a suite of simulations to test their composite effect. These simulations consist of 10%, 50%, and 90% binary characters each, where mutation rates range from 0.1 to 1 (with 0.1 increment). Due to the unexpected result of the 90% binary character scenario (see Results), I added an additional simulation with 90% ternary characters and 10% 7-state characters. I also performed an additional simulation where the proportion of binary characters ranged from 0% to 100% (with 10% increment), and with mutation rate fixed to 0.5. The mutation rate for this simulation is relatively high to reflect the typical morphological matrices, which almost always have much higher rates than molecular matrices (also see Discussion).

Next, I performed a suite of simulations which I referred to as ‘false-space’ (FS) simulations. I assigned a multistate model to matrices consisting of solely binary characters or solely ternary characters to test if SSM alone can affect reconstruction precision. This was accomplished by adding characters that are invariant across all tips, each for one possible state. I tested the effect of treating binary characters as 7-state characters, treating ternary characters as 7-state characters, and treating ternary characters as 8-state characters.

Finally, I performed an additional FS simulation with 100 32-taxa trees, each with 100 simulated matrices (10,000 matrices in total). The mutation rate for this in-depth analysis was expanded to a range between 0.1 to 2 to capture potential patterns unobserved in other simulations. I also measured weighted Robinson-Foulds distance (wRFD; Robinson and Foulds 1981), a distance measurement that accounts for branch length precision, in addition to RFD, PID, and CID. For this simulation, I included MP analysis along with the maximum likelihood analyses to look for potential correlations. I recorded not only the distances between the simulated trees and the reference trees, but also the distances among trees of different reconstruction methods. Besides distance measurements for other simulations, I performed three sets of permutation tests to better quantify the relative performance of different methods, two for testing the correlation between SSM and MP (Supplementary Tables S1 and S2), and the other for the performance between SSA and SSM (Supplementary Table S3). The comparisons were made for each mutation rate with 100,000 permutations.

### Empirical data evaluation

To illustrate how the findings apply to empirical datasets, I analyzed the morphological matrices of Amniota (focusing on Pan-Reptilia) from Bever et al. (2015) and Testudinidae (i.e., tortoises) from Torres et al. (2024) that represent typical morphological datasets. The first matrix contains exclusively fossil taxa with 85% binary characters, whereas the second includes extant taxa and has 72% binary characters. Both matrices contain characters with up to four states. Phylogenetic analyses were performed using RAxML version 8.2.10 under different state space models, including SSA, SSM, and a range of fixed-state (FS) models from five to ten states (FS5–FS10). This range was selected because SSM is effectively equivalent to an FS model with four states, and ten is the typical upper limit for numeric state codings in morphological matrices.

Character partitions for SSA were generated using MorphoParse (https://github.com/ej91016/MorphoParse), a command-line interface tool developed as part of this study in Python 3. MorphoParse accepts morphological data in commonly used formats (FASTA, PHYLIP, NEXUS, and TNT) and produces partition files compatible with RAxML (Stamatakis 2014), RAxML-NG (Kozlov et al. 2019), and IQ-TREE (Nguyen et al. 2015). It also outputs a cleaned dataset in which problematic syntax, such as special characters and polymorphisms that could lead to runtime errors, is removed. MorphoParse can thus also be used as a general-purpose tool for parsing morphological matrices and extracting only the information necessary for model-based phylogenetic inference.

In addition, strict consensus maximum parsimony (MP) trees were generated using PAUP* under the same settings for simulated data, with the rearrangement limit increased to 100,000,000 to accommodate data complexity. Pairwise Clustering Information Distance (CID) values were then calculated across the reconstructed trees. For empirical data, where the true tree is unknown, CID values were normalized against the theoretical maximum for a given tree size and topology (Smith 2020).

Description of taxonomic changes is focused on lineages of interest in the original studies. For the amniote matrix, these are the lineages with extant representative; for the tortoise matrix, it is the newly described Tablazo tortoise, a close relative of the Galapagos tortoises. These comparisons are intended to provide illustrative examples of effects that SSM can make rather than a comprehensive description of tree variations, which lies beyond the scope of this study.

## Results

### Data simulation and tree reconstruction

The control setup in which both partitions consist of binary characters results in minimal differences in tree topology and thus almost no difference in tree distance (Fig. 1a; see Supplementary Figs. S1 and S2 for PID and RFD results for all scenarios). The few cases where tree topologies are non-identical likely resulted from the final gamma optimization process, given that the recorded likelihood values are identical prior to that process. The optimization process causes more than branch length differences when the node involved is effectively a polytomy. As expected, the average tree distance between the simulated trees and reconstructed trees increases with mutation rate.

**Figure 1.**
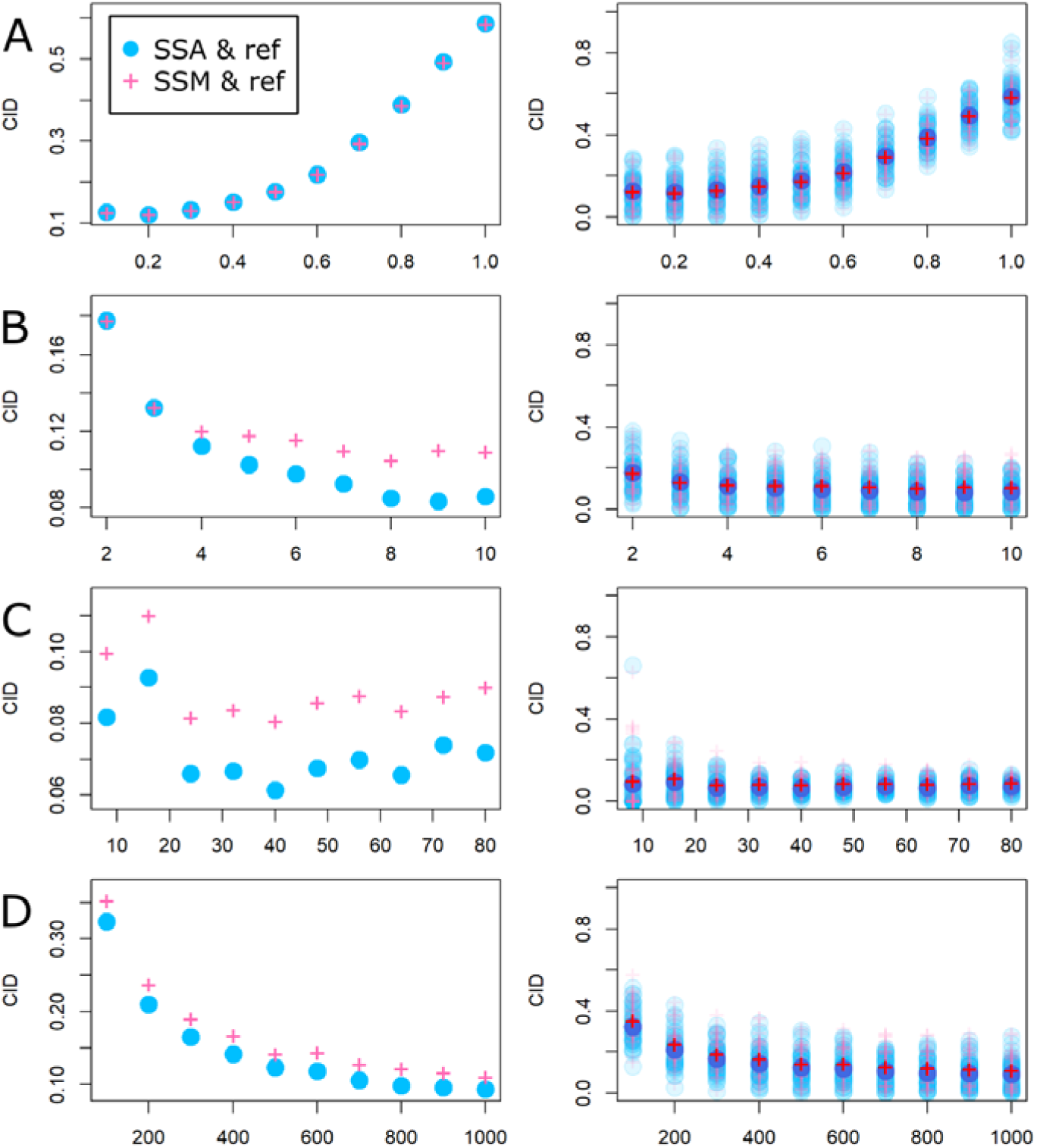
Clustering information distances (CIDs) for the control simulation and simulations with differences in ‘sizes.’ A) Control simulation; x-axis is mutation rate measured as expected number of substitutions per site per unit time. B) State space size (of the multi-state characters) difference; x-axis is state space size. C) Tree size difference; x axis is the number of tips. D) Matrix size difference; x-axis is the number of characters. Refer to Table 1 for other parameter values. Plots on the left show the average distance to the reference tree (ref) for each setup, whereas plots on the right show the distance for each simulated matrix (with the average emphasized).

When a matrix consists of both binary characters and multistate characters, the effect of SSM can be readily observed. A state space of four or greater results in a clear difference between average tree distances, where SSA outperforms SSM (i.e., have shorter average tree distances; Fig. 1b). The disparity between the two setups widens as the state space size of multistate characters increases. However, it stabilizes as the size approaches ten, as reversal becomes so unlikely that the simulation effectively applied an infinite state space model.

The tree size has no significant effect on tree distance except for RFD, and SSA consistently outperforms SSM (Fig. 1c). The 8-taxa and 16-taxa trees have higher average tree distances and variances compared to trees with more taxa. However, the absolute difference in distance is small and likely not biologically meaningful. This pattern arises because tree distance has a lower bound of zero, causing the average to be skewed toward higher values, as observed when the raw data is plotted (Fig. 1c; Supplementary Fig. S2c). However, the relatively small difference in absolute scale suggests that the results from the 16-taxa simulations remain broadly applicable. The RFD increases linearly as the tree size increases, contrasting with the consistent pattern of CID (Supplementary Figs. S1c and S2c). This contrast exemplifies well the issue of using RFD to compare trees consisting of different numbers of taxa. On the other hand, the size of the character matrix has a greater impact on tree distances, with an exponential decrease of CID as it increases (Fig. 1d). Specifically, doubling the size of a character matrix reduces CID by more than 30% when the matrix size is less than 400. I observed no clear trend between the two setups and matrix size, with the exception that SSA in all cases performed at least as well as SSM.

As the proportion of binary characters in a matrix increases, so does the tree distance, but the relative performance of SSA and SSM depends on the mutation rate (Fig. 2). When the matrices consist of 10% binary characters (Fig. 2a), minimum average CIDs are 0.06909 for SSA and 0.07030 for SSM, both of which were recorded when mutation rate equals 0.3. The maximum average CIDs are 0.10615 for SSA and 0.11223 for SSM when mutation rate equals 1. Similarly, when the proportion of binary characters increases to 50% (Fig. 2b), a mutation rate of 0.2 results in the minimum average CIDs of 0.08182 for SSA and 0.09303 for SSM. A mutation rate of 1 results in the maximum average CIDs of 0.13567 for SSA and 0.16112 for SSM. Whether it is 10% or 50% binary characters, SSA consistently outperforms SSM, but the magnitude of the difference in CIDs is greater when 50% of the characters are binary.

**Figure 2.**
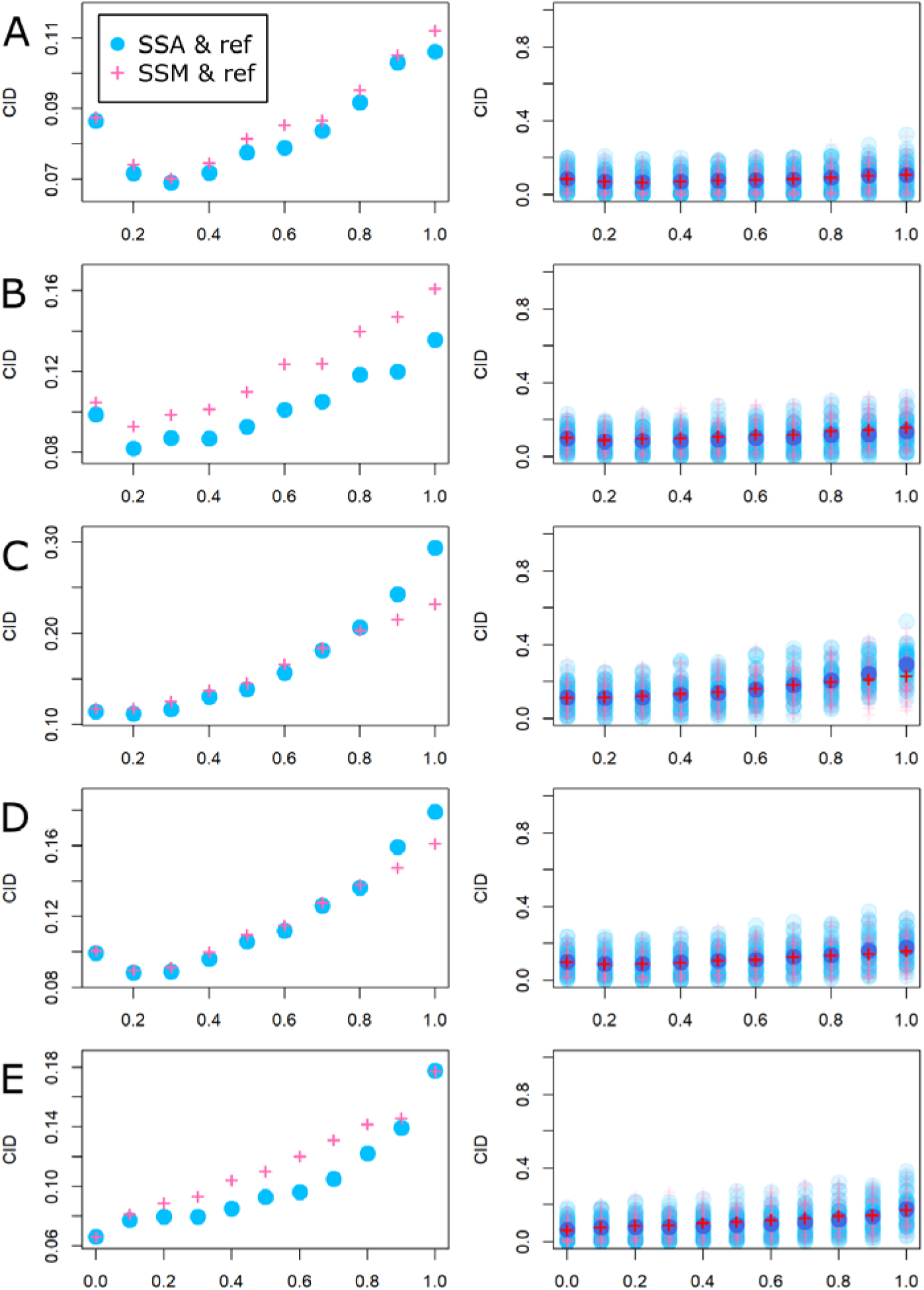
Clustering information distances (CIDs) for simulations with differences in misspecified character proportion and mutation rate. A) Rate & proportion a: 10% binary characters + 90% 7-state characters. B) Rate & proportion b: 50% binary characters + 50% 7-state characters. C) Rate & proportion c: 90% binary characters + 10% 7-state characters. D) Rate & proportion d: 90% ternary characters + 10% 7-state characters. E) Binary characters proportion difference. All x-axes are mutation rate measured as expected number of substitutions per site per unit time except for E, where x-axis is the proportion of binary characters. Refer to Table 1 for other parameter values. Plots on the left show the average distance for each setup, whereas plots on the right show the distance for each simulated matrix (with the average emphasized).

Surprisingly, a matrix of 90% binary characters diverges from this trend (Fig. 2c). Under this scenario, the minimum average CIDS are 0.11246 when the mutation rate equals 0.2 for SSA and 0.11833 when mutation rate equals 0.1 for SSM. The maximum average CIDs occurred when mutation rate equals 1 for both setups. However, SSA has a greater average CID (0.29403) compared to that of SSM (0.23289). In fact, while SSA still outperforms SSM when mutation rate is low, SSM consistently outperforms SSA when mutation rate is greater than 0.7. This implies that a misspecified model produces a more precise estimation in terms of tree topology. The same pattern emerges in simulation with 90% ternary characters (Fig. 2d), although in this scenario SSM only outperforms SSA when the mutation rate is greater than 0.8, and the difference in distance is smaller.

When the mutation rate is fixed to 0.5, SSA results in shorter average CIDs than SSM except when the percentage of binary characters is either 0 or 100 (Fig. 2e), both scenarios that involve no SSM. For both setups, average CIDs increase roughly linearly with the increased proportion of binary characters. A much greater increase in CID is observed from 90% to 100% binary characters.

For the FS scenario with binary characters, SSM starts to outperform SSA when mutation rate is greater than 0.4 (Fig. 3a). The difference between the two setups increases as the mutation rate increases, and the average CID of SSA exceeds 0.5 when mutation rate equals one. Such a large distance is unobserved in any other simulation of this study. For the FS scenario with ternary characters, SSM also outperforms SSA when the mutation rate is high (Fig. 3b, c). However, the discrepancy between SSA and SSM is greatly reduced, and the maximum average CID for SSA is less than 2.2 when mutation rate equals one. Furthermore, in both cases SSA slightly outperforms SSM when mutation rate is in between 0.3 to 0.5. The choice between 7-state or 8-state ternary character modeling has negligible impact.

**Figure 3.**
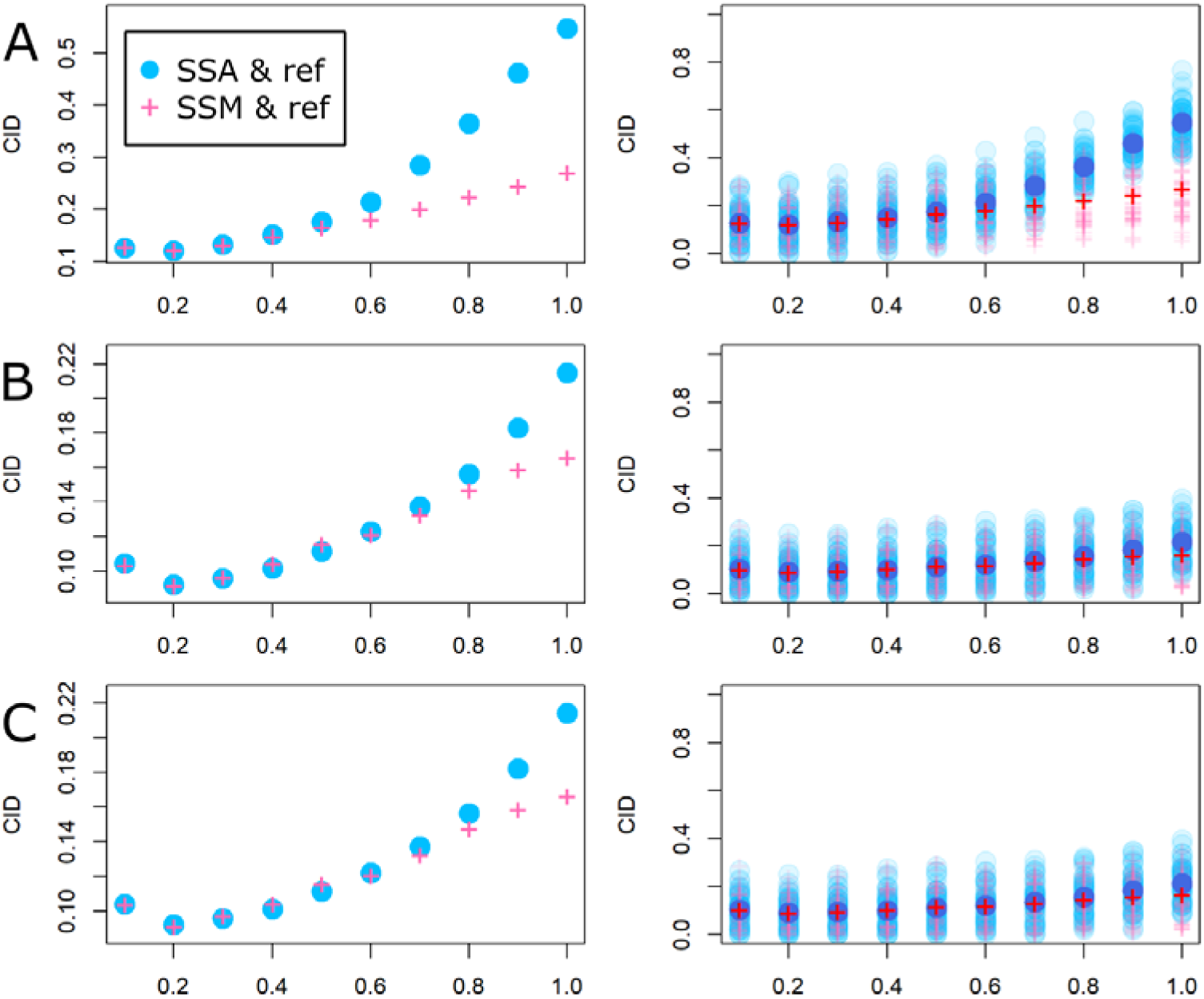
Clustering information distances for false-space simulations. A) Binary characters modeled as 7-state characters. B) Ternary characters modeled as 7-state characters. C) Ternary characters modeled as 8-state characters. All x-axes are mutation rate measured as the expected number of substitutions per site per unit time. Refer to Table 1 for other parameter values. Plots on the left show the average distance to the reference tree (ref) for each setup, whereas plots on the right show the distance for each simulated matrix (with the average emphasized).

Finally, the 32-taxa FS analysis with 10,000 matrices produces a similar pattern to the other FS analysis, where SSM increasing outperformed SSA (Fig. 4). While SSA outperforms SSM with p-value<0.05 when mutation rate ranges from 0.3 to 0.5, SSM consistently outperforms SSA with p-value <0.00001 when mutation rate is greater than 0.7. However, instead of an exponential increase, as observed when mutation rate is less than one, the surge of tree distance (specifically for SSA) slows down as mutation rate increases further. When wRFD is employed instead of the distance measurements that ignore branch length, SSA outperforms SSM when mutation rate is less than one but is eventually outperformed by SSM as mutation rate increases (Fig. 5).

**Figure 4.**
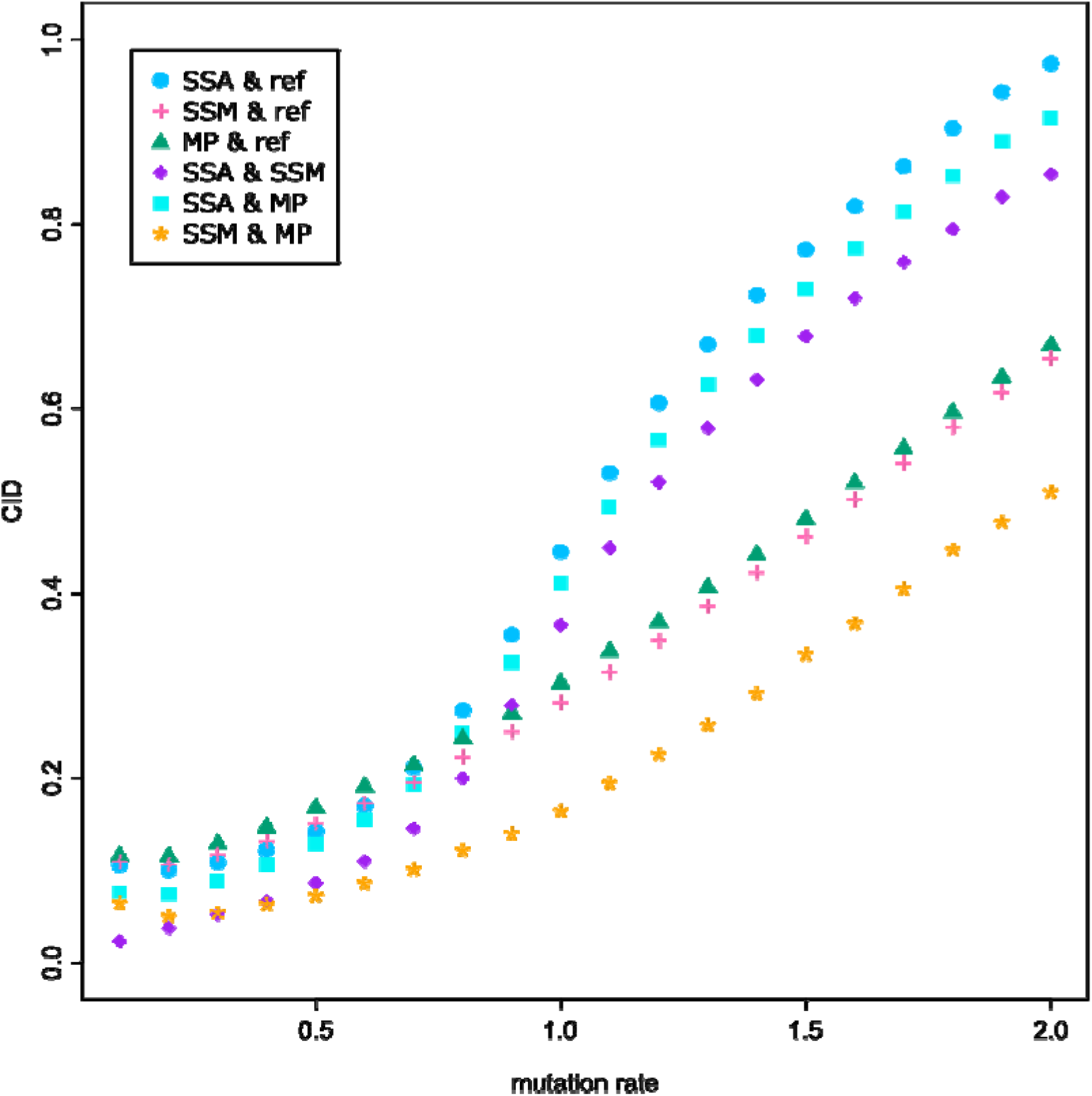
Clustering information distances (CIDs) for 32-taxa false-space simulation, including maximum parsimony and pairwise comparison. Pairwise CIDs between state space misspecification (SSM), state space-aware (SSA), and maximum parsimony (MP) reconstructions are included in addition to distances to the reference tree (ref). Mutation rate is measured as the expected number of substitutions per site per unit time.

**Figure 5.**
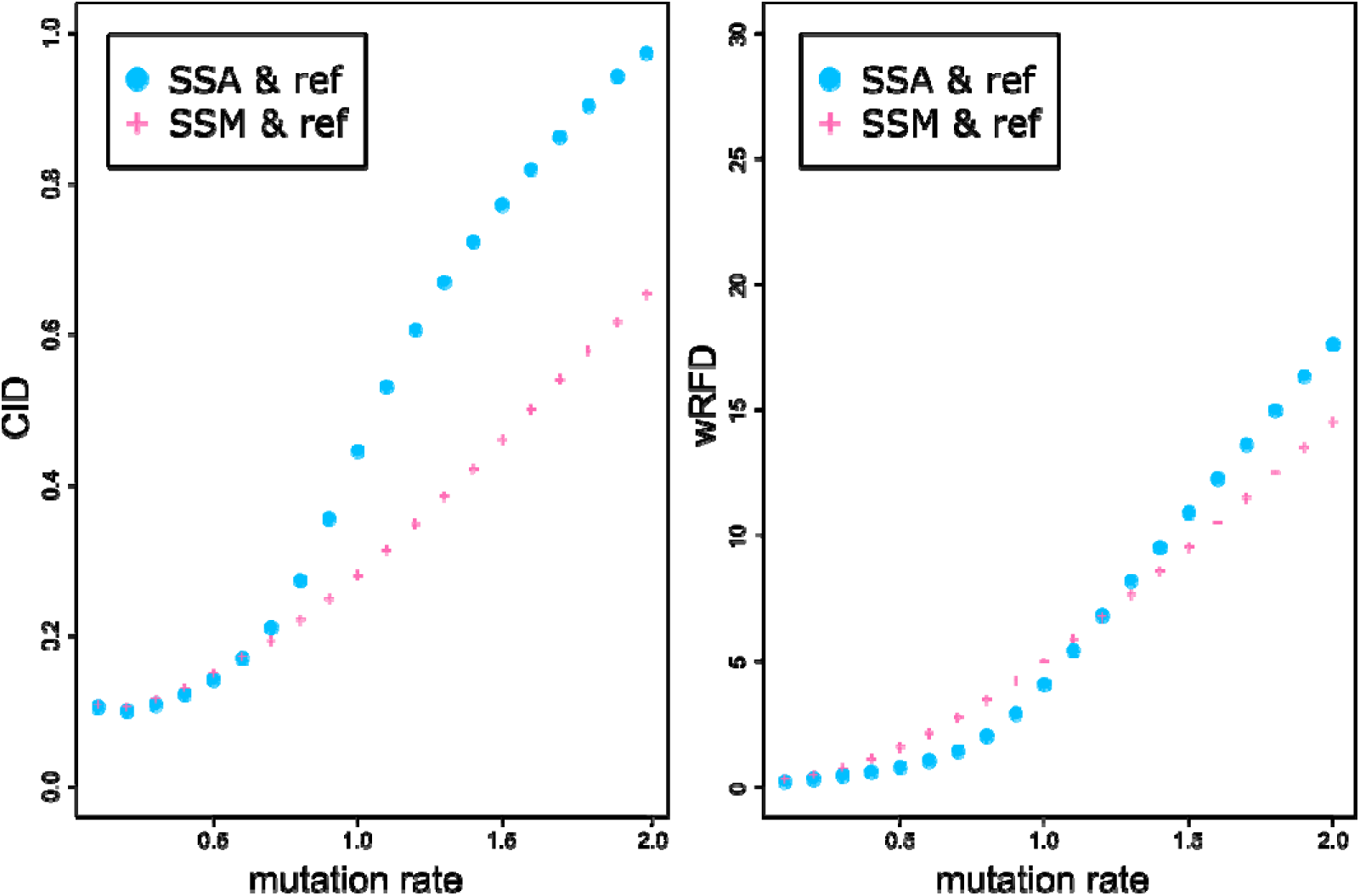
Comparison of clustering information distance (CID) and weighted Robinson-Foulds distance (wRFD) with false-space simulation. **d.** The tree set analyzed are identical for both plots. Mutation rate is measured as the expected number of substitutions per site per unit time. CID and wRFD values are not directly comparable as they have different units. However, SSA reconstruct trees more similar to the reference tree (ref) than SSM did under a greater range of mutation rate when measured with wRFD.

Comparison of tree distance between different reconstruction methods suggests complex dynamics. In general, reconstructed trees are more similar to each other than to the reference tree at low to moderate mutation rates (<0.7; Fig. 4). As the mutation rate increased, however, SSA became the outlier. Interestingly, MP consistently recovers trees with a similar level of precision (but always lower) to SSM. In fact, MP trees are not only consistently closer to SSM trees than SSA trees (p<0.00001; Supplementary Table S2), but also more similar to SSM trees (p-value < 0.00001; Supplementary Table S1) than either are to the reference trees. In contrast, SSA trees are more similar to SSM trees than SSM trees are to MP trees at low mutation rates (<0.4; Supplementary Table S3), but SSM and MP trees become closer to each other than to the SSA trees as the mutation rate further increases.

### Phylogeny with empirical data

The pairwise CID patterns are similar for both matrices (Table 2). MP trees show greater similarity to SSM and FS trees than to SSA trees. This relationship mirrors FS simulation patterns observed when mutation rates are below 0.5, though the actual CID values more closely match those seen at mutation rates around 1. For both matrices, FS reconstructions with comparable state space sizes yield similar topologies, reflecting their shared substitution model framework. However, the two matrices differ in that in the amniote matrix, the MP tree is more similar to the SSM tree than to the SSA tree, whereas for the tortoise matrix, the MP tree differs substantially from all likelihood-based trees.

**Table 2.** Pairwise clustering information distances (CID) of the empirical phylogenies. In addition to maximum parsimony (MP), state-space-aware setup (SSA), and state-space-misspecified setup (SSM), FS*n* represents a false-space (FS) simulation with a state space size of *n*. CID values are rounded to the third decimal places.

| Bever et al. 2015 | MP | SSA | SSM | FS5 | FS6 | FS7 | FS8 | FS9 |
| --- | --- | --- | --- | --- | --- | --- | --- | --- |
| SSA | 0.312 |  |  |  |  |  |  |  |
| SSM(FS4) | 0.209 | 0.210 |  |  |  |  |  |  |
| FS5 | 0.209 | 0.210 | 0.007 |  |  |  |  |  |
| FS6 | 0.203 | 0.201 | 0.009 | 0.016 |  |  |  |  |
| FS7 | 0.203 | 0.201 | 0.009 | 0.016 | 0.000 |  |  |  |
| FS8 | 0.203 | 0.201 | 0.009 | 0.016 | 0.000 | 0.000 |  |  |
| FS9 | 0.199 | 0.195 | 0.024 | 0.031 | 0.015 | 0.015 | 0.015 |  |
| FS10 | 0.199 | 0.195 | 0.024 | 0.031 | 0.015 | 0.015 | 0.015 | 0 |

| Torres et al. 2024 | MP | SSA | SSM | FS5 | FS6 | FS7 | FS8 | FS9 |
| --- | --- | --- | --- | --- | --- | --- | --- | --- |
| SSA | 0.610 |  |  |  |  |  |  |  |
| SSM(FS4) | 0.571 | 0.235 |  |  |  |  |  |  |
| FS5 | 0.571 | 0.228 | 0.032 |  |  |  |  |  |
| FS6 | 0.560 | 0.221 | 0.042 | 0.009 |  |  |  |  |
| FS7 | 0.559 | 0.213 | 0.028 | 0.023 | 0.014 |  |  |  |
| FS8 | 0.564 | 0.272 | 0.143 | 0.136 | 0.127 | 0.121 |  |  |
| FS9 | 0.560 | 0.221 | 0.048 | 0.043 | 0.034 | 0.020 | 0.101 |  |
| FS10 | 0.561 | 0.228 | 0.062 | 0.030 | 0.020 | 0.034 | 0.106 | 0.014 |

In the amniote matrix analysis, the SSA tree positioned Testudines (using *Odontochelys* and *Proganochelys* as references) basally as sister to Millerettidae. This topology recovered *Claudiosaurus germaini* as a stem ’Sauria’ (a hypothetical archosaur-lepidosaur clade) and *Acerosodontosaurus piveteaui* as a stem archosaur (Fig. 6). These results contrast with those from the SSM tree, which instead placed *Claudiosaurus germaini* as stem Reptilia and *Acerosodontosaurus piveteaui* as stem Testudines — a configuration congruent with both the MP tree and all FS trees. All maximum likelihood reconstructions consistently supported an archosaur-lepidosaur clade that excluded testudines, aligning with most morphological studies but contrasting with molecular studies that typically recover monophyletic Archelosauria (the turtle-archosaur clade). The MP tree exhibited a distinct topology, with Archosaur representing the basal divergence of Reptilia.

**Figure 6.**
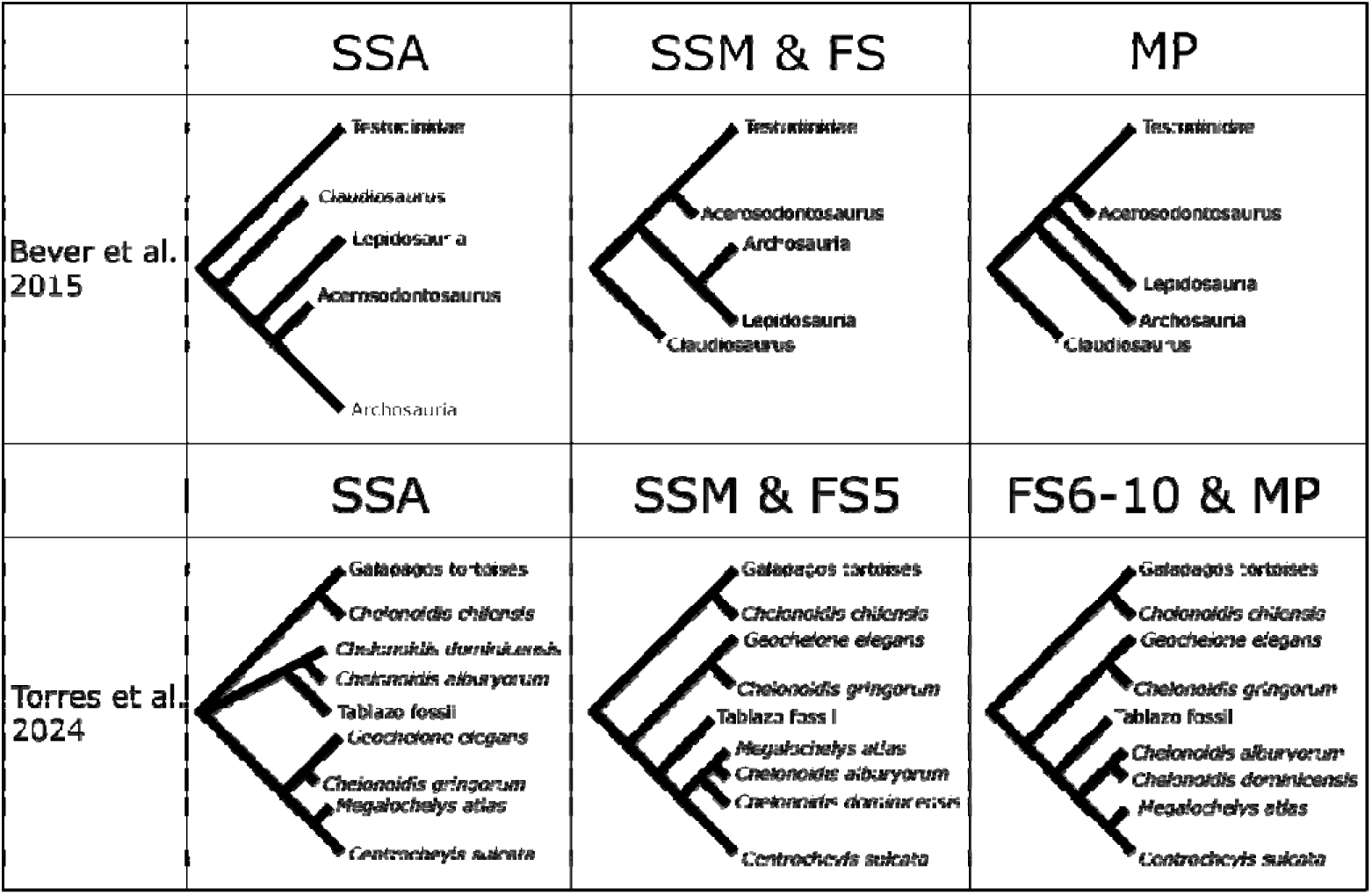
Phylogenetic reconstructions with empirical matrices. The trees summarize the variation of phylogenetic relationships among selected lineages, based upon focus of the original studies. Methods include state-space-aware (SSA) setup, state-space-misspecified (SSM) setup, false-space (FS) setup, and maximum parsimony (MP). FS represents a false-space (FS) simulation with a state space size of. The trees are mostly cladograms, except that fossil taxa are illustrated with shorter branches.

As for the tortoise matrix, the SSA tree recovered the Tablazo tortoise as sister to the extinct Caribbean tortoises *Chelonoidis alburyorum* and *Chelonoidis dominicensis*. In contrast, the SSM tree, the MP tree, and all FS trees recover the Tablazo tortoise as sister to a four-taxa clade that includes the Caribbean tortoises, *Megalochelys atlas*, and *Centrochelys sulcata*. However, the SSM tree and the FS5 tree did not recover the Caribbean tortoises as sister taxa.

## Discussion

### Phylogenetic parameters and tree distances

My simulations show that tree and matrix parameters have clear impacts on reconstruction precision, regardless of the metric of tree distance. Binary characters with high mutation rates are less ideal for phylogenetic reconstruction, as they are more prone to signal saturation (Dávalos 2014; Parins-Fukuchi 2018). The simulations also show that a mere 10 percent of multistate characters can have a major effect on the topology. While this implies inclusion of multistate characters can improve reconstruction precision, it also means they can be problematic when poorly defined.

In addition, I found a strong negative correlation between matrix size and tree distance. In this simulation, the proportion of binary characters and the tree height is on the lower side of typical proportions and branch lengths in empirical morphological phylogenies (e.g., Lewis 2001; Lee and Worthy 2011; Bever et al. 2015). But even when information is abundant compared to empirical datasets, on average RFD is improved by 1 split per 100 characters until the matrix size reaches 500. This means that even if all characters evolve in a pattern that strictly follows the species tree, it is still challenging to recover the underlying phylogeny when most characters have a small state space. To compensate, morphologists often apply additional criteria to define characters and their states (Sereno 2007; Wagner 2014). As biological constructs with evolutionary histories, all characters are, on some level, composite (Balanoff et al. 2018), and thus a matrix with 100 characters can theoretically represent hundreds of more characters. In other words, especially for multistate characters, the phylogenetic signal each character contains can be higher than appeared.

On the other hand, the observation that average tree distance strongly converges to a non-zero value after a matrix size of 500 suggests that many of the tree conflicts and polytomies populating the systematic literature are unlikely to be resolved by simply adding more characters. For speciation events that involve very short branches, resolving the order of cladogenesis may never be a realistic outcome. Even the conception of rapidly successive, distinct speciation events may be problematic given our increasing understanding of incomplete lineage sorting and hybridization (Joly et al. 2009; Meng and Kubatko 2009).

### SSA versus SSM

My simulations demonstrate that SSM plays a significant and potentially damaging role in tree reconstruction. In general, SSM results in greater tree distance between the reference and reconstructed trees. The magnitude of this distance depends on several factors, the most noticeable being differences between state space size, mutation rate, and proportion of misspecified (i.e., binary and ternary) characters. The former factor is rather trivial given that the greater the size of the assigned state space is for misspecified characters, the less appropriate the model becomes. This erosion of ‘appropriateness’ is nonlinear since the state space size is inversely proportional to the instantaneous substitution rate in the Q matrix, and consequently the calculation of likelihood values. However, increasing the state space size also means that multistate characters can carry more information, thus improving reconstruction precision. The joint effect is that the impact of SSM on reconstruction precision (i.e., differences in tree distance between SSA and SSM) is most significant when the state space size difference is moderate.

Mutation rates affect reconstruction by means of saturation or branch length. The simulations do not show a uniform relationship between mutation rate and the relative performances of SSA and SSM; rather, they magnify the trend of the relative performance. For example, when 50% of the characters are binary, SSA increasingly outperforms SSM as mutation rate increases. When all characters are treated as multistate, SSM increasingly outperforms SSA as mutation rate increases. Mutation, as a primary generator of biological variation, is thus a source of both homology and homoplasy. While the former provides meaningful structure to phylogenetic reconstruction, the latter can create noise and impair it. It is thus challenging and perhaps misleading to summarize the effects of mutation rate as unidirectional.

### Wrong model, “better” result

The counterintuitive discovery that a misspecified model situationally leads to more precise reconstruction merits at least a preliminary explanatory hypothesis. Of note, such paradoxical results occur with high mutation rate and a high proportion of binary characters, which coincidentally resembles the structure of many empirical datasets. This phenomenon is thus not only counterintuitive but also highly relevant to empirical studies that use morphological data.

That morphological matrices typically show high mutation rates and high proportions of binary characters results from a combination of biological and artificial factors. Morphological phylogenies tend to have much greater tree length and average branch length than molecular phylogenies due to how characters are defined and a phenomenon called ascertainment bias (Lewis 2001; Wright and Hillis 2014). Despite the similarity shared between molecular and morphological phylogenetics, there is a fundamental difference between the two lies in the process of data collection. Whereas molecular data are usually included in phylogenetic reconstruction regardless of distribution pattern, the establishment of morphological characters is more complicated. Individual characters are usually included with the intention to provide some phylogenetic information, an inherent qualification is that these characters bear some variation. Consequently, a typical morphological matrix would have a higher estimated mutation rate than a molecular matrix, which contains many more invariant sites.

Similarly, morphological characters are predominantly binary due to two clear advantages for the working scientist. First, a binary character requires a simple decision: with the presence/absence characters that comprise many binary characters, a specimen either expresses the described character state or it does not. This simplicity not only increases efficiency in scoring characters, but also fits well with an adaptive narrative, that a character was gained or lost by a lineage through some adaptive process. This is especially relevant given that many characters are initially identified as synapomorphies and autapomorphies of specific taxa (Assis and Rieppel 2011). Second, although gray areas might exist between two states in a binary character, the size of the state space can be fixed at two with certainty. This fixation of state space size at two enjoys the benefits of model simplicity, since the order of the states is irrelevant (which is especially applicable to maximum parsimony) and it requires minimal effort to exhaust the observed variation for the given character.

Nevertheless, there is a tradeoff for such convenience. Simplifying character variation means a reduction of information and increases the probability of saturation and convergence. While neither of these phenomena are necessarily problematic for phylogenetic reconstruction (and model-based approaches explicitly acknowledge them to some extent), extensive saturation and convergence can still lead to significant loss of phylogenetic signal (Brinkmann et al. 2005). We thus arrive at an unpleasant situation where in addition to their relatively small size compared to molecular matrices, current morphological matrices are also more susceptible to phylogenetic ‘noise’ due to the limited information discrete morphological characters typically contain.

But why is the noise that is introduced by binary characters reduced by SSM, a clearly incorrect model? This counterintuitive effect of SSM on tree topology is demonstrable with a classic four-taxa phylogeny. We can build a simple scenario where taxon A and taxon B exhibit state0 of a binary character k, while taxon C and taxon D exhibit state 1. Figure 7 shows two of the possible topologies: these balanced trees are identical with branch lengths equal to 0.5, except Ta proposes a homologous origin of the states, whereas Tb proposes a homoplastic origin.

**Figure 7.**
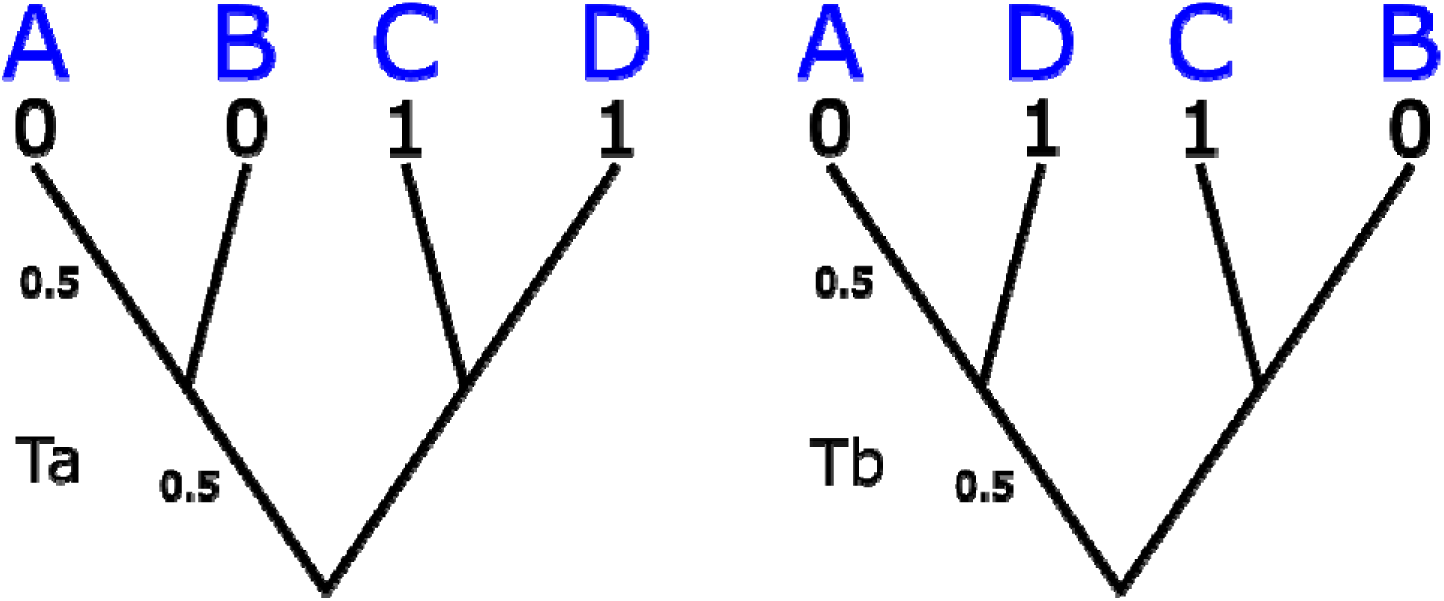
Two possible 4-taxa phylograms. All branches are assumed equal length and measured as expected number of mutations per site per unit time. Ta requires minimally 1 mutation event, whereas Tb requires minimally 2 mutation events.

Under a 2-state MK model, the ln-likelihood (lnL) values for the two topologies with respect to character k are:

On the other hand, if character k is treated as a 7-state character, then under a 7-state MK model:

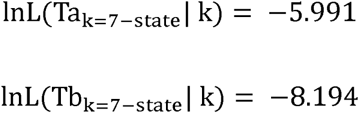

Despite both showing a preference toward homologous origin of the state (Ta), the magnitude of support is much stronger under a 7-state model. This is because under a time reversible model, the greater the number of possible states is, the less likely a primary homology is homoplasy in reality. Suppose the ln-likelihood for the two trees with respect to the rest of the matrix (M) differs by one such that:

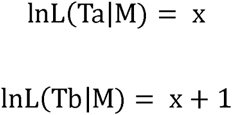

Then under a 2-state model, Tb is the more likely hypothesis:

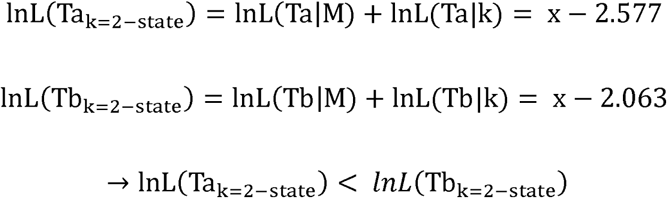

In contrast, under a 7-state model, Ta is the more likely hypothesis:

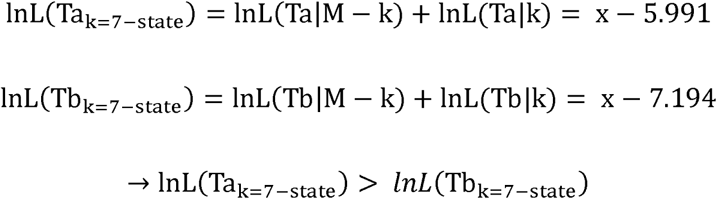

The difference in inferred signal strength is thus important when other characters do not show clear alignment with a specific tree hypothesis, as a SSM model can create a bias by overemphasizing a subset of characters.

Since this bias is site-specific, not tree-specific, its overall effect depends on the number of characters supporting each topology. If we assume a positive correlation (even just a loose one) between the number of inferred homoplasy and distance to the true tree, one that is made explicit in maximum parsimony, SSM would lead to an overall bias toward trees that are closer to the true tree. Additionally, as the bias is more likely to overturn the topological hypothesis when the likelihood for each tree is similar, it would have greater impact when data contain more noise (as introduced by relatively less informative binary characters), as observed in the simulations. On the contrary, one would expect that this bias is reduced as the true state space increases. Indeed, the simulations show that the paradoxical phenomenon is reduced or even reverted when binary characters are replaced by ternary characters (Fig. 2c,d).

Although the bias SSM causes may appear advantageous in terms of topology when data is limited, it introduces undeserved preference. SSM overemphasizes binary characters, potentially altering tree hypotheses that multistate characters (if present) would support. Furthermore, trade-offs exist with the increase of topological precision, as SSM underestimates branch length. An intuition is that the longer the branches are, the less likely the characters have yet explored the entirety of the state space. This can also be demonstrated by the four-taxa example above. Consider Ta with identical branch length x, the likelihood is maximized at x=0.275 under a 2-state MK model and x=0.218 under a 7-state MK model. As a result, when evaluated using wRFD, the relative performance of SSM declines significantly compared to methods that do not incorporate branch length. Although branch lengths measured in mutation rate are rarely discussed in morphological studies, they are parameters directly involved in likelihood and posterior probability calculation, with potential impacts on efforts to incorporate time into cladograms using dating. The biased branch length estimates produced by SSM are thus highly relevant to model-based phylogenetic methods. Altogether, the success of SSM under certain circumstances (i.e., high proportion of binary character and high mutation rate) comes with critical trade-offs: it may obscure the phylogenetic signals provided by multistate characters while also producing branch lengths that are underestimated.

### SSM and morphological phylogenetics

While the observed patterns are mostly drawn from simulations that one may argue to be over-simplistic, the behavior can be explained by a clear statistical framework. Analyses with empirical matrices confirmed the general applicability of these interpretations. Notably, both the amniote and tortoise matrices exhibited the predicted sensitivity to state space specification, with topological differences between SSM and SSA reconstructions mirroring simulation patterns. The results demonstrate high sensitivity of certain branches to minor model adjustments — an instability often overlooked in phylogenetic studies. Although this study focused on ML, the implications extend to Bayesian frameworks, which similarly rely on likelihood calculations for morphological data. This relationship becomes especially clear when considering that significant discrepancy between prior assumption and the likelihood of the data pattern should be considered problematic (Lemmon et al. 2004). The applicability to Bayesian inference is particularly relevant given the growing importance of morphological data in tip-dating analysis, where fossils provide critical time calibration (Arcila et al. 2015).

Whereas molecular phylogenetics has moved away from MP for tree reconstruction, it is still a standard practice in morphological phylogenetics. The similarity between reconstructed SSM and MP trees are thus particularly applicable to morphological studies. Trees reconstructed with different methods can be more similar to each other than to the reference tree due to homoplastic patterns in the matrices, which introduce noise that biases topologies away from the underlying phylogeny. However, the observation that SSM trees are more similar to MP trees than to SSA trees (except when mutation rates are low) suggests a methodological consistency beyond mere matrix identity.

As discussed in the introduction, standard MP will always select the same topologies as maximum likelihood (ML) under NCM model, assuming uniform state spaces across characters. In effect, even without making this assumption explicit, unweighted MP behaves as though state space variation is ignored. This situation can be understood in one of two ways: either as an acceptable artifact of MP’s simplicity, or as a misalignment that demands correction.

In the next section, I provide a formal derivation showing how to reconcile MP with a correctly specified NCM model that accounts for variation in character state space. The resulting solution is a SSA weighting scheme in which each character is weighed by the natural logarithm of its state space size. This approach ensures that MP will select the same topologies as ML under a variable-state-space NCM model. SSA weights can be implemented automatically using MorphoParse, which outputs a compatible weighting block for PAUP* and TNT.

### State-space-aware weighting reconcile MP and the NCM model

Theorem 5 of Tuffley and Steel (1997) shows that maximum likelihood estimate using no common mechanism model (NCM) will select the same tree(s) as maximum parsimony. Specifically, the maximum likelihood conditioned on topology T is given by

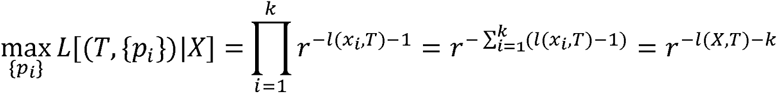

Where {pi} is the set of mutation probability on each branch for character *i, x_i_* is the state pattern of character *i*, *l*(*x_i_*,*T*) is the minimal inferred number of mutations given *x_i_* and topology T, and r is the number of possible states. The expression 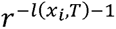 arises because each of the *l*(*x_i_*,*T*) branches with an inferred mutation contributes a probability of 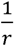, and the probability of root state contributes to an additional factor 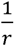. It follows that the likelihood is maximized when *l*(*X*, *T*), the number of steps required for a topology, is minimized. However, an assumption here is that the size of state space is identical for all characters. This assumption is generally valid for molecular sequences, but rarely held in morphological datasets, where characters often vary substantially in the number of possible states.

A maximum likelihood estimate that acknowledges state space variation, and therefore appropriate for morphological matrices should bear the form of

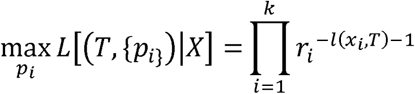

Where *r_i_* represents the number of possible states for character *i*, and can expand to

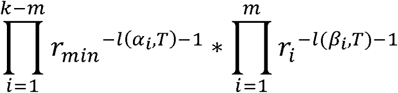

Where, {*α_i_*} are the characters with the minimal state space and {*β_i_*} are the characters with non-minimal state space. It follows that

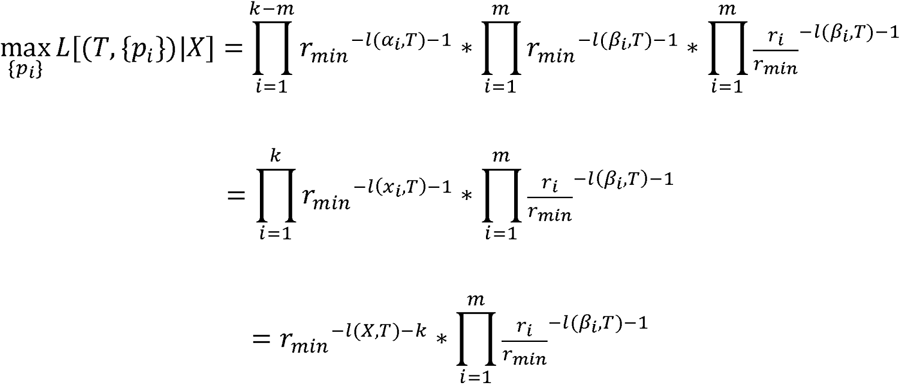

Let T(s) denote the most parsimonious reconstruction and *l(β_i_,T(s))* the steps required for character *β_i_*, with non-minimal state space. Let T(s’) denote an alternative topology with *l*(*X,T*(*s’*)) = *l*(*X,T*(*s*)) + *t* where *t ∊ N*. For maximum likelihood estimate with NCM to always choose the same topology as maximum parsimony, it needs to be such that

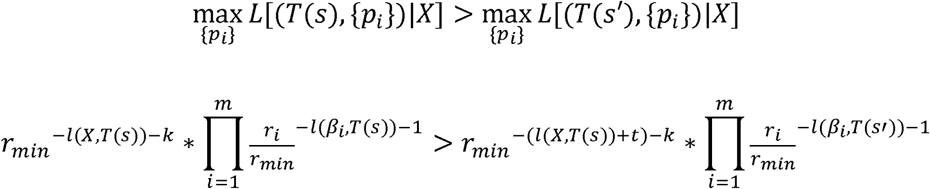

This can be simplified to

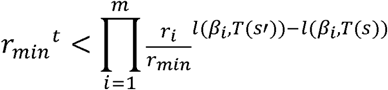

and with log transformation,

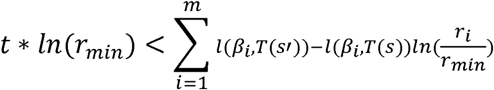

Since *r ∊ N and t ∊ N*, the inequality is trivially impossible if *r_i_* = *r_min_ ∀_i_* or *l*(*β_i_*, *T*(*s’*)) − *l*(*β_i_*, *T*(*s*) ≤ 0 *∀_i_*. The first scenario would mean equal state space size for all characters, which implies to state space variation. The second scenario would mean that topology T is the most parsimonious for every character with non-minimal state space size.

Unless under the aforementioned scenarios, there is no guarantee that the two methods select the same topology. For example, consider a topology that differs from the most parsimonious topology by exactly one step for a character c with non-minimal state space. The above inequality can be simplified as

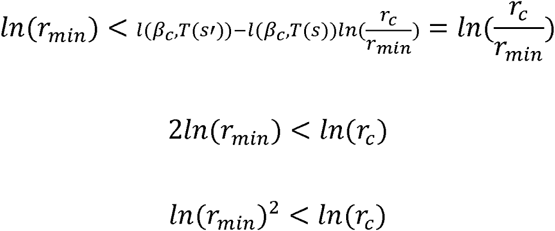

Let *r_min_* = 2, which means there is at least a binary character in a matrix, then maximum likelihood estimate with NCM will prefer a topology with an extra state so long as character c has 5 or more possible states.

A full mathematical enumeration of the conditions under which NCM and maximum parsimony select the same or different topologies can be an interesting problem, but for phylogenetics a more pressing question is the adjustment required to maintain the consistency for the two approaches. If we were to keep the choice of maximum parsimony, a solution is to add an assumption to NCM — that all characters are assumed to have the same state space size. The modeled state space size can be arbitrary large to ensure that all observed characters state can be properly allocated. Such assumption can seem unrealistic for most characters but has been described and discussed in the past (e.g., Cuthill 2015), and is technically applied when practicing standard parsimony if considered under a model-based framework.

Alternatively, the practice of maximum parsimony can be adjusted to always select the same tree(s) as maximum likelihood estimate with NCM by introducing a weight scheme. Specifically, when characters are weighted by ln (*r*) where *r* represents the number of states for each character such that 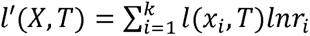 is the weighted parsimony score, then

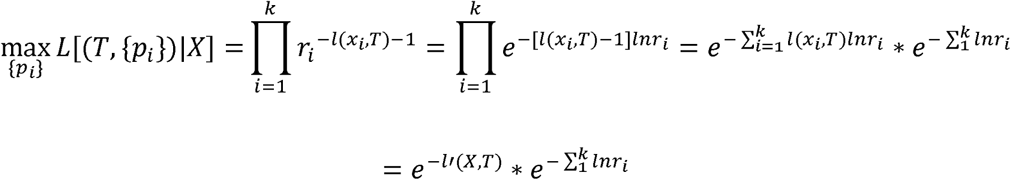

Since the second term is a constant that does not depend on the tree topology, it is trivial that 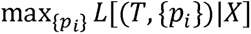 is maximized when *l*’(*X, T*) is minimized. This means one step for a character with larger state space carry more weight than one step for a character with smaller state space. The rationale is that given equal substitution rate, the larger the state space the less likely there would be a reversal. In other words, primary homologies observed for characters with large state space are more likely to be true homology and therefore granted more weight.

It is worth noting that standard maximum parsimony always selecting the same tree(s) as a maximum likelihood method that ignores state space variation does not mean parsimony itself disregards state space (Sober 2004). Rather, as shown here, unweighted MP behaves like a method that fails to account for state space variation. While the NCM model has been cited in defense of MP for both its topological agreement and philosophical alignment, this connection breaks down when characters vary in their number of states. Applying weights to characters may appear to violate MP’s appeal to simplicity, but uniform weighting is itself a model — one that implicitly assumes equal state space across all characters. Given the SSM-like behavior of unweighted MP, such an assumption is problematic. If state space has any meaningful role in morphological data, and MP is to be reconciled within a probabilistic framework, then weighting by ln (*r*) is not just a theoretical option but a necessary adjustment.

A full exploration of the SSA weighting scheme’s empirical performance is beyond the scope of this study, as it would require testing under a range of generating models, minimally including both common mechanism and no common mechanism scenarios. However, the weighting scheme introduced here ensures statistical consistency with the NCM framework, and it places MP on a firmer theoretical footing, comparable to model-based methods, despite concerns about the NCM model’s overparameterization (Holder et al. 2010).

### Limitations and applications

A key strength of simulation-based studies is that parameters (e.g., tree topology, mutation rate, and state space) are known with certainty, allowing a direct exploration of their relationships with reconstruction outcomes. In contrast, empirical datasets offer only estimates of such parameters, often confounded by hidden processes. This distinction is apparent in the two empirical matrices analyzed here. Both reflect the simulation-predicted similarity between SSM and MP trees, but only the amniote matrix shows a stronger resemblance between SSM and MP than between SSM and SSA. Moreover, the degree of similarity is weaker than that seen in the 32-taxa FS simulations. This discrepancy likely reflects complex biological and methodological processes in real datasets that are absent from the simulation framework. As such, caution is warranted when extrapolating simulation results to empirical analyses. A broader evaluation of SSM effects in empirical studies may require a larger-scale meta-analysis across diverse morphological matrices, particularly to investigate factors like fossil sampling, character coding strategies, and binary character prevalence shape sensitivity to state space assumptions.

Returning to the simulation framework, the reference pure-birth trees used in this study involved only two parameters: the birth (diversification) rate and the tree size. They are equivalent to extant phylogenies where taxa are randomly sampled, but unlike many morphological phylogenies in that they do not contain fossil representatives. Fossils have the potential to break down long branches, thus reducing the various effects associated with saturation and increasing reconstruction precision. In fact, this could be the reason why SSM tree and SSA tree are more similar to each other than to the MP tree for the two empirical datasets in this study despite the effects of a high mutation rate discussed above. One interesting area of future research lies in investigating the consequences of replacing pure-birth trees with birth-death or fossilized birth-death trees (Heath et al. 2014).

Similarly, data simulated in this study evolved under simple models where most parameters are fixed both temporally and spatially. The effects of greater variation in character-state spaces, in addition to rate heterogeneity and clock models are among many potential avenues of further study. Although it would be intriguing to pursue a comprehensive study on the effects of SSM, exhausting all combinations of potential parameters is unlikely to be accomplished within a reasonable timeframe. Two more practical approaches are either to increase the flexibility of parameters while only drawing conclusions based on general patterns thence observed, or to restrict parameter values and draw conclusions with specific comparisons. In this study, I chose the latter and in-so-doing established a foundation for studying the relationship between SSM and several critical parameters. I also showed throughout the study that binary characters play a leading role in the potential effects of SSM on phylogenetics.

Despite the limitations of simulation, the results of this study can provide valuable insights on dealing with SSM in empirical datasets. Given the complex effect of SSM on precision in tree reconstruction, a meaningful solution might be just as complex. However, there are several practices that can be adopted with relatively minor effort to decrease potential effects of SSM. For existing matrices, one can simply run an extra analysis where FS is introduced alongside SSA. I recommend a state space size of 10 to maximize the chance of topological convergence without introducing non-numeric state for a standard matrix. Additional analyses of different FS sizes can also collectively provide a more robust comparison regarding the uncertainty of topology, especially given the paradoxical effect of SSM under certain circumstances. A caveat here is that the absolute branch lengths should not be interpreted, as they will be underestimated in most cases.

In contrast to ML, MP has the opposite problem where state space needs to be recognized. Imposing explicit assumptions on what are poorly understood and likely complex dynamics of morphological characters can be intimidating, however as mentioned the standard MP is not as assumption-free as it might appear. The SSA weighting scheme where characters are weighed by ln(r) simply adjust the ‘MP model’ such that character state space can be properly handled.

Another approach is to explicitly incorporate uncertainty into a model. For a small number of characters, this can be done by estimating the size of state space for each character either through maximum likelihood or Bayesian inference. On the other hand, when dealing with a large number of characters or when direct estimation of state space size is challenging, this can be done using similar approaches to the use of a gamma distribution to describe rate heterogeneity across sites (Yang 1996).

Finally, an approach that targets the fundamental issue is to reduce the number of binary characters in a matrix. This includes removal of highly homoplastic binary characters, but more importantly the establishment of more multistate characters. Transforming a binary character into a multistate character requires extra work but allows a more genuine portrayal of true biological variation. The use of continuous characters is another interesting alternative, as it would theoretically maximize the information each character can hold. The incorporation of continuous characters in phylogenetics has received increased attention recently (Parlin-Fukuchi 2018, Tagliacollo et al. 2025), but remains an exploratory area of research.

The complexity of morphological data, from character definition to state delimitation has led to the dominance of molecular phylogeny, where models are well-defined and data are abundant. However, molecular sequences are not always available, particularly in deep-time studies where the vast expanse of evolutionary history has been obscured by extinction and the only remaining evidence for that history is found in fossils. Ancient DNA is an exciting possibility for recent lineages (Pääbo et al. 2004; Kjær et al. 2022), but the relatively rapid degradation of DNA molecules leaves little hope for application to nodes deeper than a couple million years (Dalén et al. 2023). In these scenarios, morphological data remains a crucial, and probably our only, source of phylogenetically informative data. In addition, morphological data are critical to the on-going pursuit of realizing a robust genotype-phenotype map (Wagner and Zhang 2011; Morris et al. 2020). Rapidly advancing technologies in both sequencing and editing of genomes represent an unprecedented opportunity for understanding how molecular information translates to functional morphological systems and the organisms they serve (Bressloff 2017; Feng et al. 2022). In the genotype/phenotype dichotomy, it is the phenotype (and especially the morphological phenotype) that has become the empirical and analytical bottleneck (Schönenberger et al. 2020). Finally, it should not be forgotten that the morphological phenotype plays a critical mechanistic role in the functioning of ecological and evolutionary systems as the most direct target of selection (Kimura and Crow 1978; Kingsolver et al. 2001). There are ample reasons, therefore, to continue and even amplify our efforts to improve precision in the phylogenetic analysis of morphological data.

## Conclusion

The quality of morphological character data is frequently discussed, especially in light of the clarity of homology statements (character definitions), the empirical basis for polymorphism levels, the extent of potential homology, as well as many other considerations. Much less attention has been directed to the quantitative specification of these characters, especially when a model-based analysis is involved. In this study, I show that state space is a neglected aspect of morphological character analysis with significant potential for shaping our perception of evolutionary history. My simulations collectively demonstrate that SSM is a condition that can interact with other parameters and affect the precision of phylogenetic reconstruction. In addition, binary characters are especially susceptible to SSM, resulting in complicated and even paradoxical behavior. It is important to note that this study does not diminish the use of binary characters altogether, but rather emphasizes the merits of more detailed documentation of character variation. This includes the circumscription of multistate characters. While we remain a long way from a full understanding of the degree to which SSM impacts tree topologies and patterns of character evolution, it is my hope that this study will encourage further investigation into of the role of state space uncertainty in phylogenetic reconstruction by means of both simulated and empirical data.

## Acknowledgements

I thank Gabriel Bever, Amy Balanoff, Jake Wilson, Will Foster, Fernando Torres, and Emma Puetz for their discussion and feedback on the manuscript. I also appreciate James Boyko’s suggestion to investigate branch length precision, which proved to be a critical factor. I acknowledge the use of ChatGPT for assistance in refining the implementation and streamlining the development of MorphoParse.

## Data Availability

All simulation scripts are available on GitHub at https://github.com/ej91016/SSM, and simulation data and results are archived in the Dryad Digital Repository (http://datadryad.org/share/k541YB5iaYxIEWxZsinyUgVfsl6jxtsJ4u40YNPdFGQ). The Python tool MorphoParse, used for matrix parsing, partitioning, and implementing the SSA weighting scheme, is available at https://github.cosm/ej91016/MorphoParse.

## Notes

https://github.com/ej91016/MorphoParse

https://github.com/ej91016/SSM

### Competing Interest Statement

The authors have declared no competing interest.

