## Supplementary for "State Space Misspecification in Morphological Phylogenetics: A Pitfall for Models and Parsimony Alike"

EJ Huang

Center for Functional Anatomy & Evolution, Johns Hopkins University School of Medicine, Baltimore, MD 21205, USA

Corresponding author: EJ Huang

**This PDF file includes:**

**Figures S1 to S2**

**Tables S1 to S3**

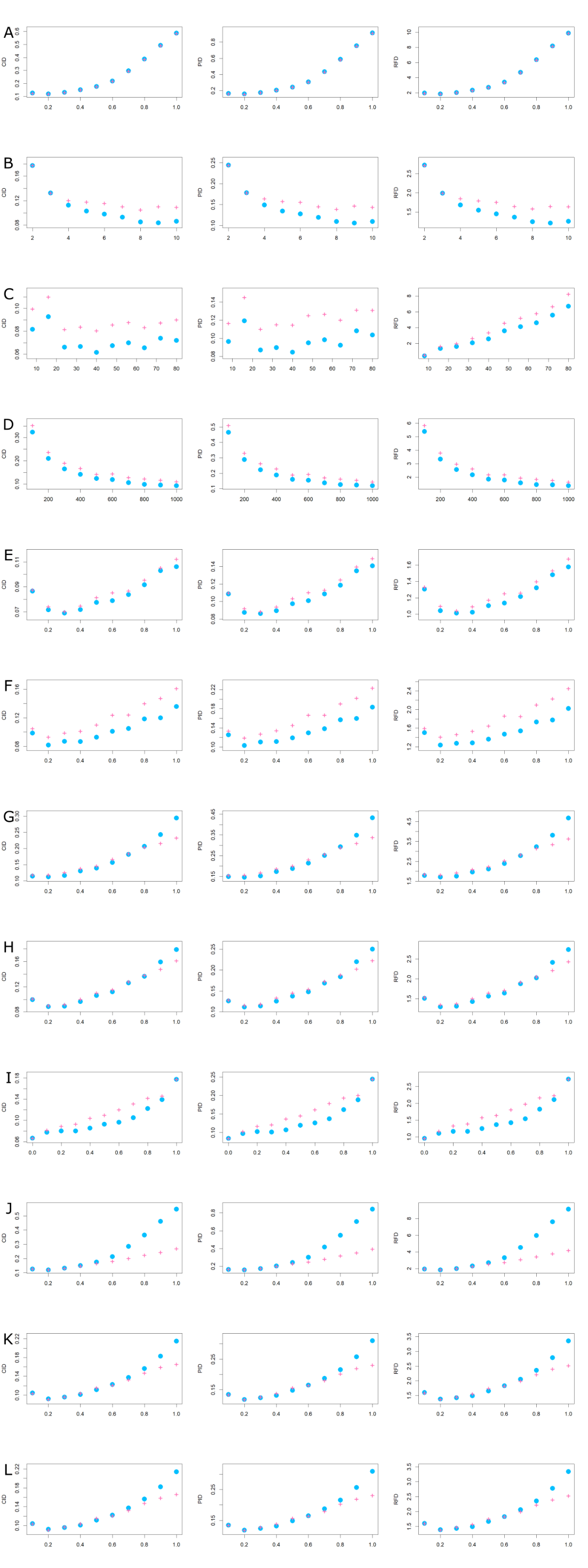

**Supplementary Figure S1. Average tree distances CIDs for all simulations.** The average lustering information distance (CID), phylogenetic information distance (PID) and Robinson-Foulds distances (RFD) between the reconstructed trees and the reference tree. All x-axes are mutation rate unless otherwise specified. A) Control simulation. B) State space size (of the multi-state characters) difference; x-axis is state space size. C) Tree size difference; x axis is number of tips. D) Matrix size difference; x-axis is number of characters. E) Rate & proportion a: 10% binary characters + 90% 7-state characters. F) Rate & proportion b: 50% binary characters + 50% 7-state characters. G) Rate & proportion c: 90% binary characters + 10% 7-state characters. H) Rate & proportion d: 90% tertiary characters + 10% 7-state characters. I) Binary characters proportion difference. K) Binary characters modeled as 7-state characters. K) Tertiary characters modeled as 7-state characters. L) Tertiary characters modeled as 8-state characters. Refer to Table 1 for parameter values. Refer to Table 1 in the main article for parameter values.

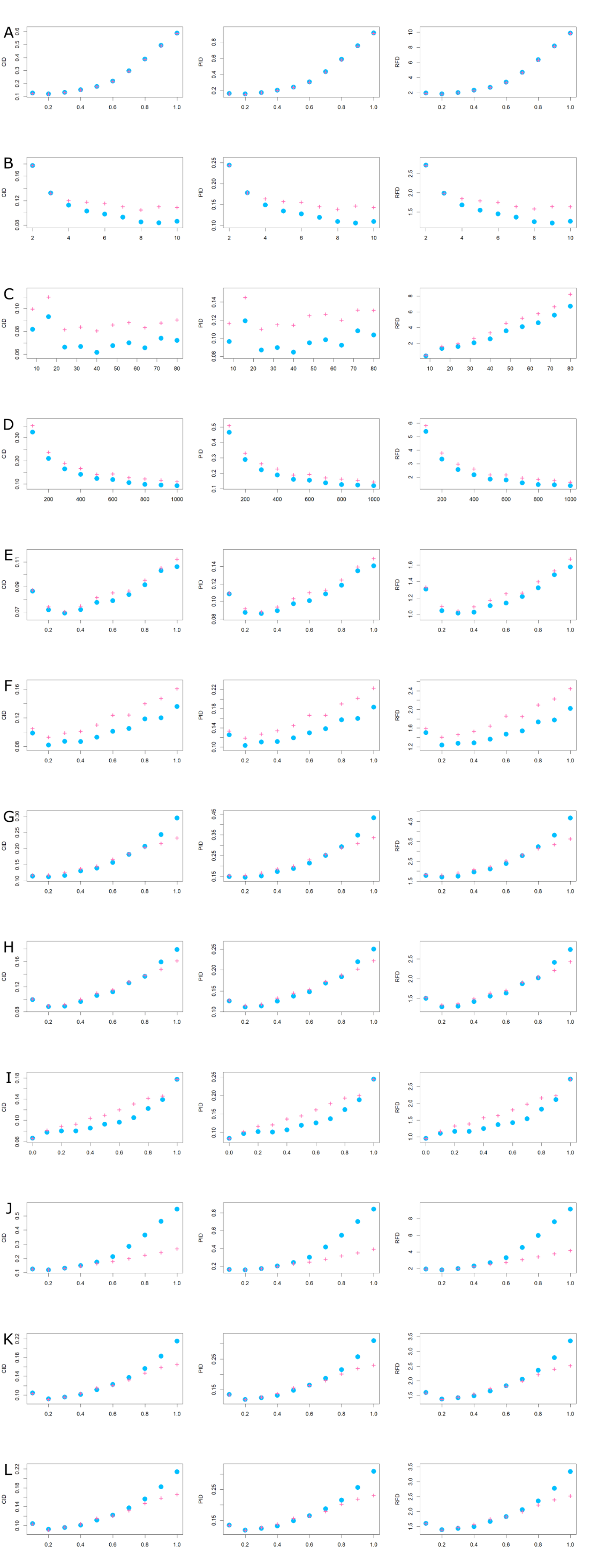

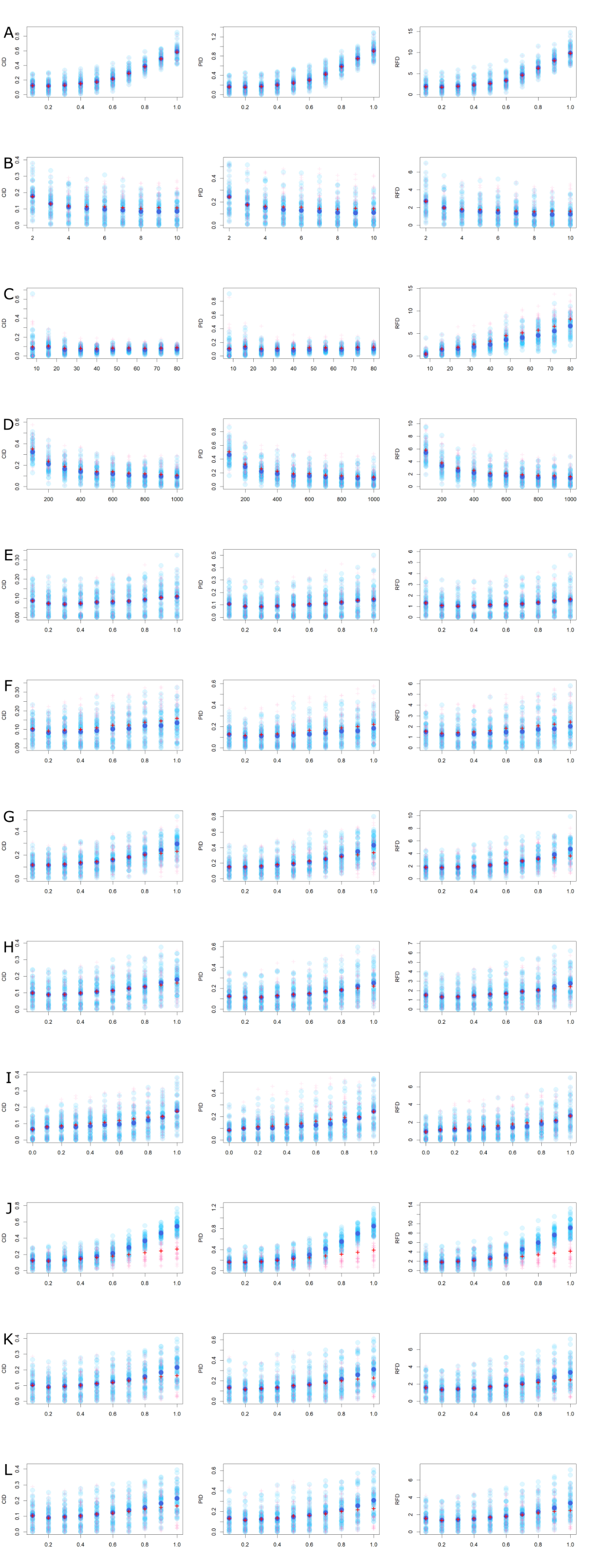

**Supplementary Figure S2. Tree distances for all simulations.** Clustering information distance (CID), phylogenetic information distance (PID) and Robinson-Foulds distances (RFD) between the reconstructed trees and the reference tree. All x-axes are mutation rate unless otherwise specified. A) Control simulation. B) State space size (of the multi-state characters) difference; x-axis is state space size. C) Tree size difference; x axis is number of tips. D) Matrix size difference; x-axis is number of characters. E) Rate & proportion a: 10% binary characters + 90% 7-state characters. F) Rate & proportion b: 50% binary characters + 50% 7-state characters. G) Rate & proportion c: 90% binary characters + 10% 7-state characters. H) Rate & proportion d: 90% tertiary characters + 10% 7-state characters. I) Binary characters proportion difference. J) Binary characters modeled as 7-state characters. K) Tertiary characters modeled as 7-state characters. L) Tertiary characters modeled as 8-state characters. Refer to Table 1 for parameter values. Refer to Table 1 in the main article for parameter values.

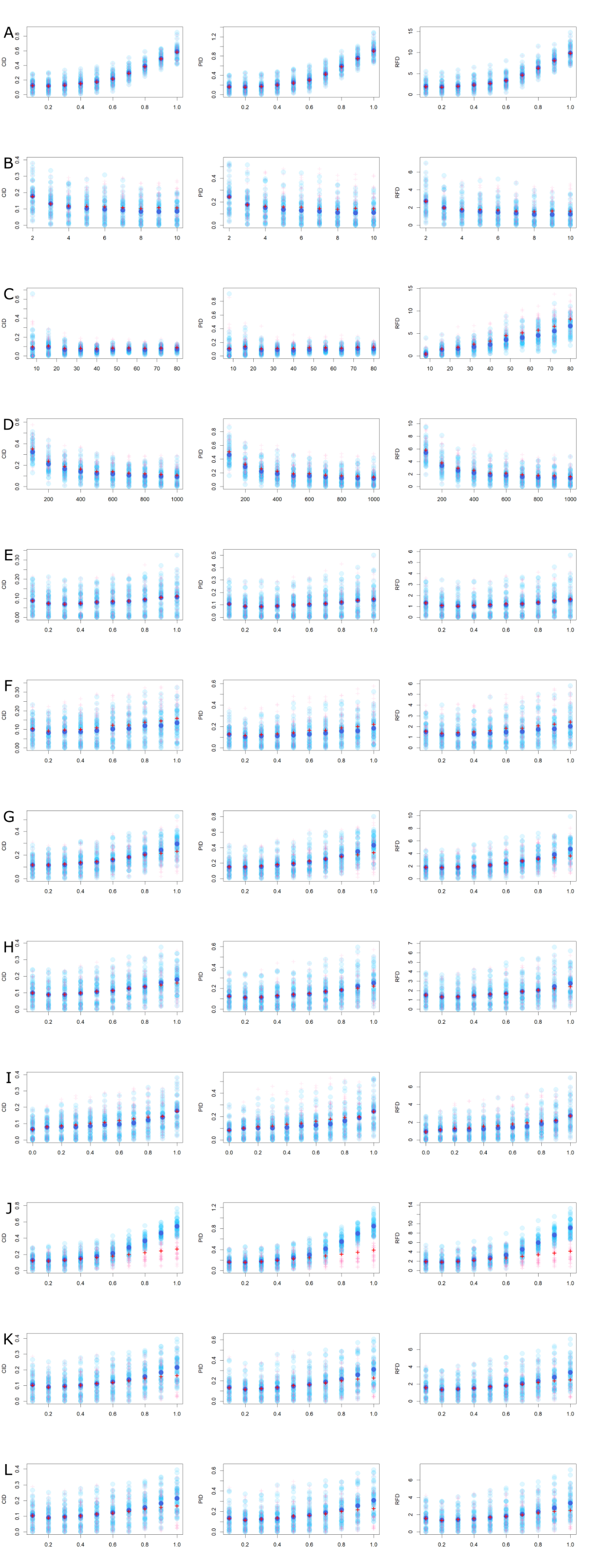

**Supplementary Table S1. Permutation Test for similarity between maximum parsimony (MP) and the model-based methods.** Results of permutation tests based on the 32-taxa false-space simulation d (FS d) assessing whether clustering information distance (CID) is smaller between the MP tree and the state-space-misspecified (SSM) tree or between the MP tree and the state-space-aware (SSA) tree. ‘closer.to.SSM’ indicates whether the MP tree is closer to the SSM tree (TRUE) or not (FALSE). The associated p-value reflects the statistical significance of the difference. Mutation rate is measured as expected number of substitutions per site per unit time.

| **mutation rate** | **closer.to.SSM** | **p-value** |
| --- | --- | --- |
| 0.1 | TRUE | <0.00001 |
| 0.2 | TRUE | <0.00001 |
| 0.3 | TRUE | <0.00001 |
| 0.4 | TRUE | <0.00001 |
| 0.5 | TRUE | <0.00001 |
| 0.6 | TRUE | <0.00001 |
| 0.7 | TRUE | <0.00001 |
| 0.8 | TRUE | <0.00001 |
| 0.9 | TRUE | <0.00001 |
| 1 | TRUE | <0.00001 |
| 1.1 | TRUE | <0.00001 |
| 1.2 | TRUE | <0.00001 |
| 1.3 | TRUE | <0.00001 |
| 1.4 | TRUE | <0.00001 |
| 1.5 | TRUE | <0.00001 |
| 1.6 | TRUE | <0.00001 |
| 1.7 | TRUE | <0.00001 |
| 1.8 | TRUE | <0.00001 |
| 1.9 | TRUE | <0.00001 |
| 2 | TRUE | <0.00001 |

**Supplementary Table S2. Permutation Test for similarity between the state-space-misspecified setup (SSM) and maximum parsimony (MP).** Results of permutation tests based on the 32-taxa false-space simulation d (FS d) assessing whether clustering information distance (CID) is smaller between the SSM tree and the reference tree or between the SSM tree and the MP tree. ‘closer.to.ref’ indicates whether the SSM tree is closer to the reference tree (TRUE) or not (FALSE). The associated p-value reflects the statistical significance of the difference. Mutation rate is measured as the expected number of substitutions per site per unit time.

| **mutation rate** | **closer.to.ref** | **p-value** |
| --- | --- | --- |
| 0.1 | FALSE | <0.00001 |
| 0.2 | FALSE | <0.00001 |
| 0.3 | FALSE | <0.00001 |
| 0.4 | FALSE | <0.00001 |
| 0.5 | FALSE | <0.00001 |
| 0.6 | FALSE | <0.00001 |
| 0.7 | FALSE | <0.00001 |
| 0.8 | FALSE | <0.00001 |
| 0.9 | FALSE | <0.00001 |
| 1 | FALSE | <0.00001 |
| 1.1 | FALSE | <0.00001 |
| 1.2 | FALSE | <0.00001 |
| 1.3 | FALSE | <0.00001 |
| 1.4 | FALSE | <0.00001 |
| 1.5 | FALSE | <0.00001 |
| 1.6 | FALSE | <0.00001 |
| 1.7 | FALSE | <0.00001 |
| 1.8 | FALSE | <0.00001 |
| 1.9 | FALSE | <0.00001 |
| 2 | FALSE | <0.00001 |

**Supplementary Table S3. Permutation Test for relative performance between state-space-aware setup (SSA) and state-space-misspecified setup (SSM).** Results of permutation tests based on the 32-taxa false-space simulation d (FS d) assessing whether clustering information distance (CID) is smaller between the SSM tree and the reference tree or between the SSM tree and the maximum parsimony tree. ‘better’ indicates the method that performs better given the mutation rate. The associated p-value reflects the statistical significance of the difference. Mutation rate is measured as the expected number of substitutions per site per unit time.

| **mutation rate** | **better** | **p-value** |
| --- | --- | --- |
| 0.1 | SSM | 0.7576 |
| 0.2 | SSA | 0.07214 |
| 0.3 | SSA | 0.00741 |
| 0.4 | SSA | 0.00377 |
| 0.5 | SSA | 0.0215 |
| 0.6 | SSM | 0.65712 |
| 0.7 | SSM | <0.00001 |
| 0.8 | SSM | <0.00001 |
| 0.9 | SSM | <0.00001 |
| 1 | SSM | <0.00001 |
| 1.1 | SSM | <0.00001 |
| 1.2 | SSM | <0.00001 |
| 1.3 | SSM | <0.00001 |
| 1.4 | SSM | <0.00001 |
| 1.5 | SSM | <0.00001 |
| 1.6 | SSM | <0.00001 |
| 1.7 | SSM | <0.00001 |
| 1.8 | SSM | <0.00001 |
| 1.9 | SSM | <0.00001 |
| 2 | SSM | <0.00001 |
